# Inspecting Morphological Features of Mosquito Wings for Identification with Image Recognition Tools

**DOI:** 10.1101/410449

**Authors:** Clinton Haarlem, Rutger Vos

**Affiliations:** Naturalis Biodiversity Center, Leiden, the Netherlands; Institute of Biology Leiden, Leiden, the Netherlands

## Abstract

Mosquitoes are important disease vectors. Different mosquito genera are associated with different diseases at varying levels of specificity. Hence, quick and low-cost methods of identification, even if relatively coarse and to genus level, will be of use in assessing risk and informing mitigation measures. Here we assess the extent to which digital photographs of mosquito wings taken with common cell phone cameras and clip-on lenses can be used to discriminate among mosquito genera when fed into image feature extraction algorithms. Our results show that genera may be distinguished on the basis of features extracted using the SURF algorithm. However, we also found that the naïve features examined here require very standardized photography and that different phone cameras have different signatures that may need to be taken into account.

## 1. Introduction

### 1.1 Mosquito Biology

The mosquito is a small winged insect that belongs to the order Diptera and the family Culicidae. Mosquitoes are a very large group of insects and the Culicidae family is a monophyletic taxon that consists of 3,490 currently recognized species grouped in 44 genera [1] (see appendix I for table). This group is not only very species rich, it is also extremely abundant and diverse. Mosquitoes occur in every part of the world except for Antarctica and they can develop in a broad range of biotic communities such as arctic tundra, boreal forests, deserts, mountains, marshes and ocean tidal zones. The greatest species diversity occurs in tropical forests [2]. Mosquitoes generally feed on plant nectar, but females of most species are parasitic and must consume blood to gain nutrients and proteins needed to produce their eggs. They can feed on a diversity of hosts, ranging from mammals and birds to reptiles, amphibians and even fish [3,4]. This parasitic behaviour has earned the mosquito the reputation of being a notorious pest. Mosquitoes have the capability to spread dangerous diseases which have made them one of the most deadly animal groups on the planet [5,6].

#### Morphology

Mosquitoes come in many shapes and sizes and can morphologically differ in numerous ways. The greatest differences are apparent when comparing certain genera and can often be distinguished with the naked eye, but differences between closely related species can be microscopically small and difficult to recognize [7,8]. Differences in morphology may be found in the wings, abdomen, female and male genitalia, the head, thorax, legs, mouth parts, scales and setae and often need expert knowledge to locate [9]. Differences between species can also be visible in different life stages of the insect. For instance, larvae may be identified by their abdominal segments, siphon, setae or teeth [10].

#### Breeding preferences

Besides morphological differences, biological factors such as longevity, habitat preference and feeding behaviour may also be different between species [11]. One clearly observable difference between the groups is breeding biology. All Diptera, and therefore all mosquitoes, go through four different life stages: egg, larva, pupa and imago (adult). Because the Culicidae group is so large, there is much variation in the life cycles of different genera and species of mosquito. Some species may have multiple broods of offspring per year and others may have only one. Some species are able to overwinter in the egg stage, whereas others overwinter either as larvae or as adults [12]. The larvae of all mosquitoes are aquatic and therefore eggs must be laid in or near bodies of water. Mosquitoes from the genus *Aedes* lay single eggs on moist substrates. These eggs can withstand a certain amount of desiccation. Mosquitoes from the genera *Anopheles, Culex* and *Mansonia* lay their eggs in water. *Anopheles,* like *Aedes*, lays eggs in single units, whereas *Culex* and *Mansonia* lay clusters off eggs. The clusters of *Mansonia* are attached on the underside of vegetation. The egg rafts of *Culex* float on the water’s surface [13]. Another difference in breeding preference is seasonality: in a study in Pakistan, larvae of the genera *Anopheles* and *Aedes* were mostly collected in July, whereas larvae of the genus *Culex* were mainly collected in September [14]. Available sunlight also plays a role in the selection of a breeding site, as some *Anopheles* species have a preference for water that is exposed to the sun [15]. *Anopheles* and *Culex* mosquitoes generally lay their eggs on the surface of permanent water pools. *Aedes* individuals tend to lay eggs in soil in areas where flooding occurs, such as flood banks, ditches and irrigated pastures. They also lay eggs in pools formed after rain or in containers filled with water [16]. Mosquitoes from the genus *Anopheles* have been thought to prefer clean, clear water to breed, whereas individuals from the genera *Culex* and *Aedes* prefer water high in organic compounds, such as found in polluted water, septic tanks and gutters [17], although it has recently been discovered that several *Anopheles* species can tolerate and breed in polluted water as well [18,19]. Different properties and parameters of water such as pH level, hardness, temperature, chemical composition and the presence of bacterial fauna may all influence which species of mosquito choose to breed in which locations. Thus, the condition and quality of water in an area directly affect the presence of specific species of mosquito.

#### Mosquito-borne disease

Mosquitoes play an important role in the transmission of disease and can be vectors for pathogenic bacteria, viruses and parasites. Mosquito borne diseases such as malaria, dengue, yellow fever and West-Nile virus cause millions of deaths a year [5,6]. Not all mosquitoes are known disease vectors, and many diseases are only carried by a specific genus or species. Table 2 displays an overview of mosquito borne diseases and their respective vectors.

**Table 1:** Mosquito borne diseases and their respective vectors.

| Disease | Vector | Source |
| --- | --- | --- |
| Chikungunya | *Aedes aegypti*, *Aedes albopictus* | [6,20] |
| Dengue | *Aedes aegypti*, to a lesser extent *Aedes albopictus* | [6,21] |
| Japanese encephalitis | *Culex* spp. | [22] |
| Malaria | *Anopheles* spp., the *gambiae* and *funestus* complexes in particular. | [23,24] |
| West-Nile | Most *Culex* spp., to a lesser extent *Aedes albocephalus* and *Mimomyia* spp. | [6,25] |
| Yellow fever | *Aedes aegypti* | [6,26] |
| Zika | *Aedes* spp. | [6,27] |
| Lymphatic filariasis | *Culex* spp., *Aedes* spp., *Anopheles* spp., *Mansonia* spp. and *Ochlerotatus* spp. | [28,29] |

**Table 2:**
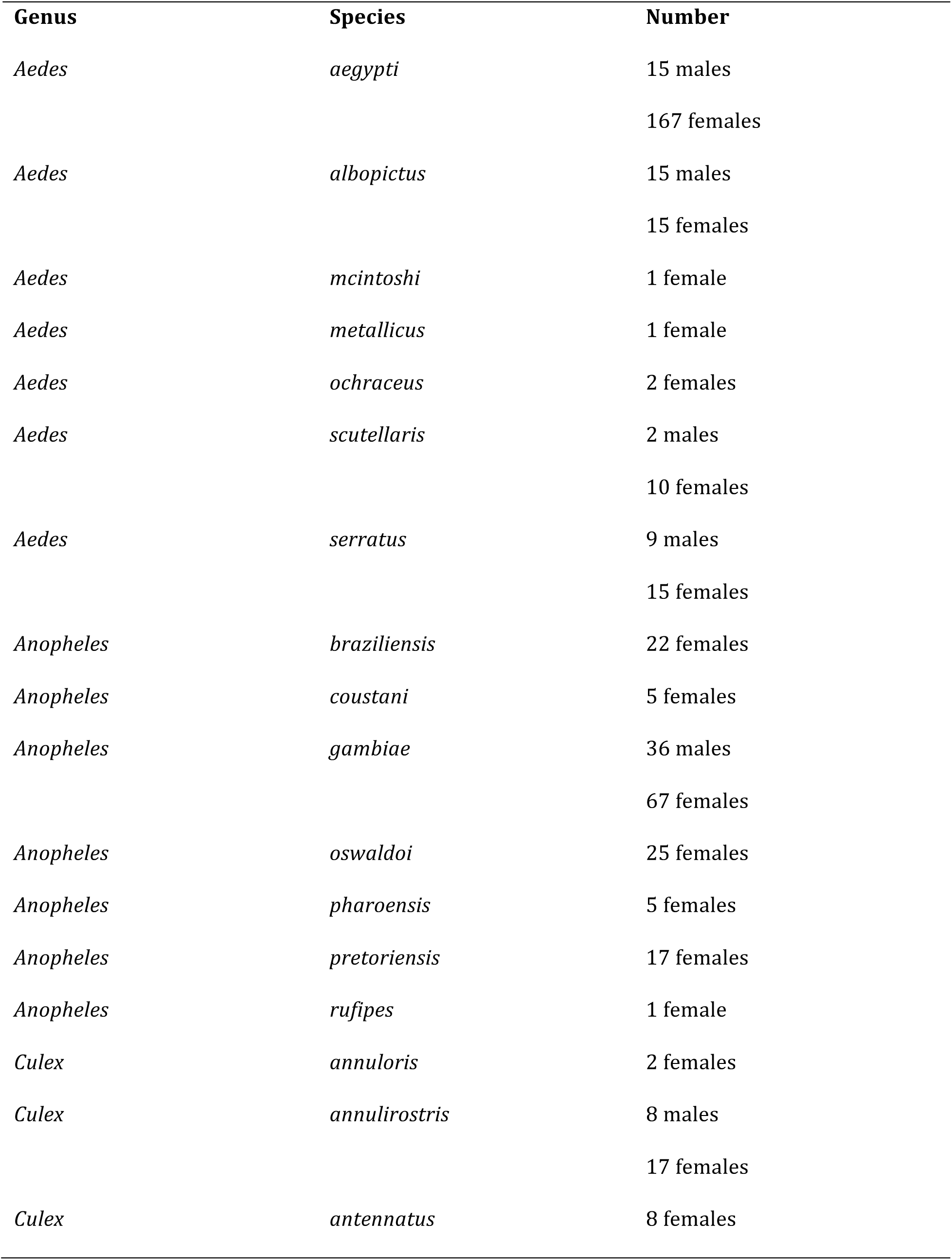
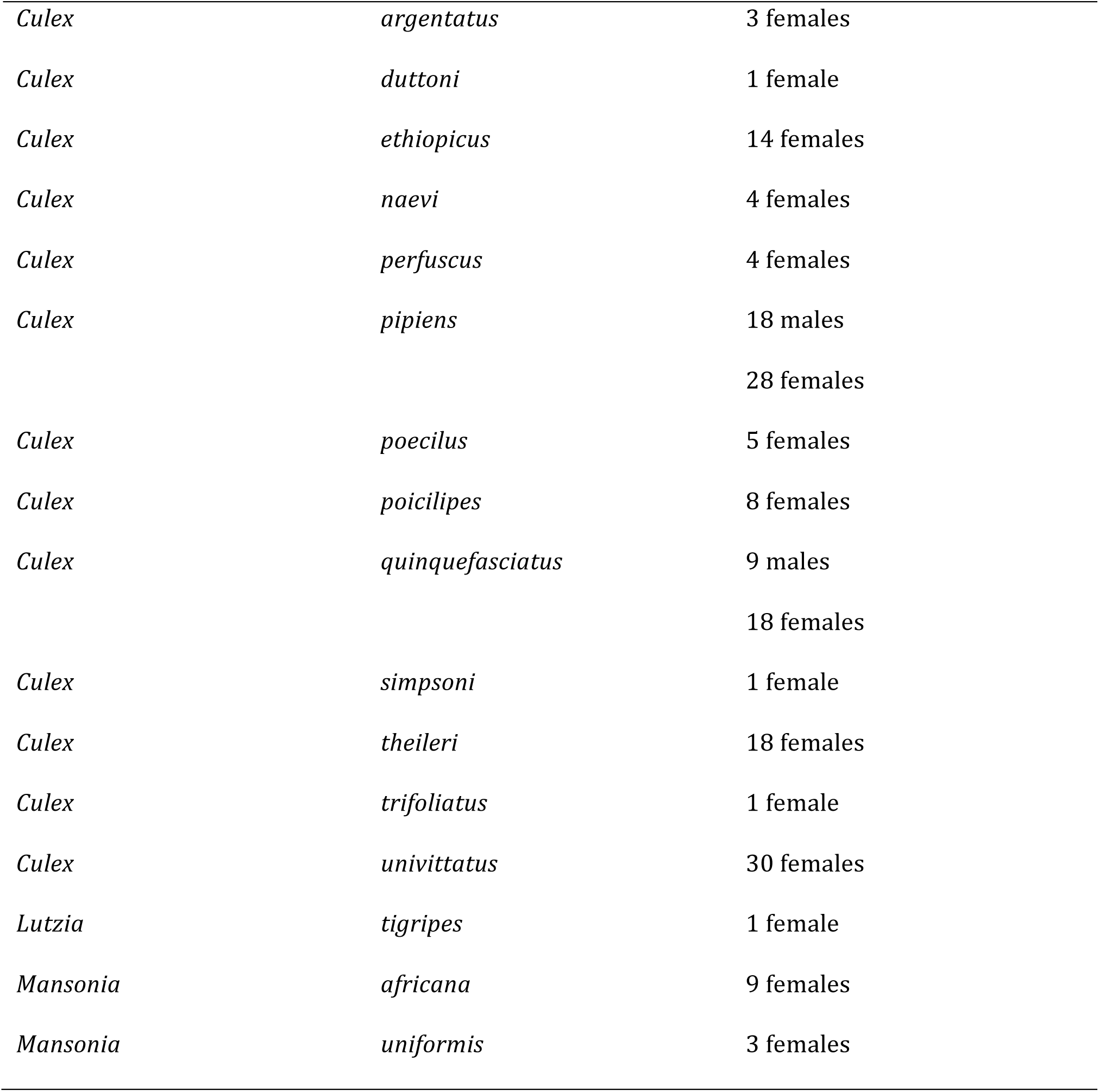
Genus, species, number and sex of the mosquitoes photographed for this project.

| Genus | Species | Number |
| --- | --- | --- |
| *Aedes* | *aegypti* | 15 males |
|  |  | 167 females |
| *Aedes* | *albopictus* | 15 males |
|  |  | 15 females |
| *Aedes* | *mcintoshi* | 1 female |
| *Aedes* | *metallicus* | 1 female |
| *Aedes* | *ochraceus* | 2 females |
| *Aedes* | *scutellaris* | 2 males |
|  |  | 10 females |
| *Aedes* | *serratus* | 9 males |
|  |  | 15 females |
| *Anopheles* | *braziliensis* | 22 females |
| *Anopheles* | *coustani* | 5 females |
| *Anopheles* | *gambiae* | 36 males |
|  |  | 67 females |
| *Anopheles* | *oswaldoi* | 25 females |
| *Anopheles* | *pharoensis* | 5 females |
| *Anopheles* | *pretoriensis* | 17 females |
| *Anopheles* | *rufipes* | 1 female |
| *Culex* | *annuloris* | 2 females |
| *Culex* | *annulirostris* | 8 males |
|  |  | 17 females |
| *Culex* | *antennatus* | 8 females |
| *Culex* | *argentatus* | 3 females |
| *Culex* | *duttoni* | 1 female |
| *Culex* | *ethiopicus* | 14 females |
| *Culex* | *naevi* | 4 females |
| *Culex* | *perfuscus* | 4 females |
| *Culex* | *pipiens* | 18 males |
|  |  | 28 females |
| *Culex* | *poecilus* | 5 females |
| *Culex* | *poicilipes* | 8 females |
| *Culex* | *quinquefasciatus* | 9 males |
|  |  | 18 females |
| *Culex* | *simpsoni* | 1 female |
| *Culex* | *theileri* | 18 females |
| *Culex* | *trifoliatus* | 1 female |
| *Culex* | *univittatus* | 30 females |
| *Lutzia* | *tigripes* | 1 female |
| *Mansonia* | *africana* | 9 females |
| *Mansonia* | *uniformis* | 3 females |

Monitoring vectors is an important part of controlling and possibly preventing these diseases. Because strains of pathogens are often carried by specific mosquito species or genera [6,30], an essential aspect of monitoring disease vectors is to keep track of the distribution and abundance of the different species in certain areas. However, in areas where mosquito-borne disease is a big problem, knowledge about the carriers is often limited [31,32,33] and mosquitoes are subsequently not properly monitored [23]. This often leaves the problematic mosquito species unidentified or their distribution and abundance uncertain. Another possible problem in monitoring mosquito distribution is that climate change may alter the range of species [34].

### 1.2 Mosquito identification

To keep track of which mosquito species may be involved in the transmission of which diseases, it is important to accurately identify these species. Because mosquito identification based on morphology is difficult, it generally requires specialist entomological and taxonomic knowledge and hard to come by identification keys. DNA barcoding may prove to be an effective alternative, but this method requires expensive equipment and specifically designed labs [35,36,37]. A method that would require none of these to identify mosquito species would greatly benefit scientific research. Not only would an identification tool that does not rely on specialist knowledge or equipment be a cheap and easy to use solution for scientists and researchers, a readily accessible identification method would also allow local communities to become more involved in mapping mosquito-borne diseases. Recruiting locals in making mosquito observations would considerably increase the amount of data collected and, by providing the users of the tool with information about the species they are mapping, would simultaneously increase local knowledge and awareness of the occurrence of particular diseases in an area.

#### Image recognition

For species with clear morphological differences, image recognition could be a valuable method for the development of an identification tool. Image recognition software is able to recognize specific colour ratios, colour distribution, the presence of patterns or unique markings, dimensions and shapes [38,39,40]. Because many species from various phyla do have distinctive morphological features, image recognition software has become a popular tool in the taxonomic community. Several applications have already been developed to identify a range of different organisms, from orchids to fruit flies to bees to spiders [41,42,43, 44, 45]. In flying insects, the wing seems to be a very distinguishing morphological feature. Some identification applications were even designed around this idea and focus mostly on geometric morphometric analyses to recognize wing shape and venation patterns. Examples include DrawWing [46], ABIS [47], DAIIS [48] and WingID [43].

#### Wing morphometrics

Mosquito wings also appear to have some distinctive features, as they have already been used to distinguish between genera [37], between species within the same genus [49,50,51], between populations of a species [52,53] and between sexes of a species [54]. Geometric morphometric analyses in these studies were carried out by comparing landmark points between wings using techniques such as Procrustes analysis. Landmarks were identified as wing vein branch points and intersections between veins or between a vein and the wing margin. All studies used 18 landmarks, except for Mondal et al. [50], in which 20 landmarks were identified (Figure 1). These publications show that although all studied mosquito species share similar landmark points, morphometric differences were clear enough to distinguish between taxonomic groups. Mosquito wings may therefore be a suitable focus for an automated identification tool based on image recognition. This study aims to test the functionality of using mosquito wings for automated identification purposes by creating an image database of mosquito wings from several different genera and species and by analysis this database using different image recognition algorithms. The end goal of the project is the creation of an identification application that can be used on smartphones and tablets. Users should ultimately be able to take a photograph using their mobile device, upload it to the app and subsequently receive information about the photographed specimen. The focus of this project lies with South-African genera/species. South-Africa is home to many species of mosquito, both harmless and disease carrying [55]. The problematic species can carry a number of different diseases and having an identification app that focuses on this region could have an impact on areas such as health and medicine, education and local welfare.

**Figure 1:**
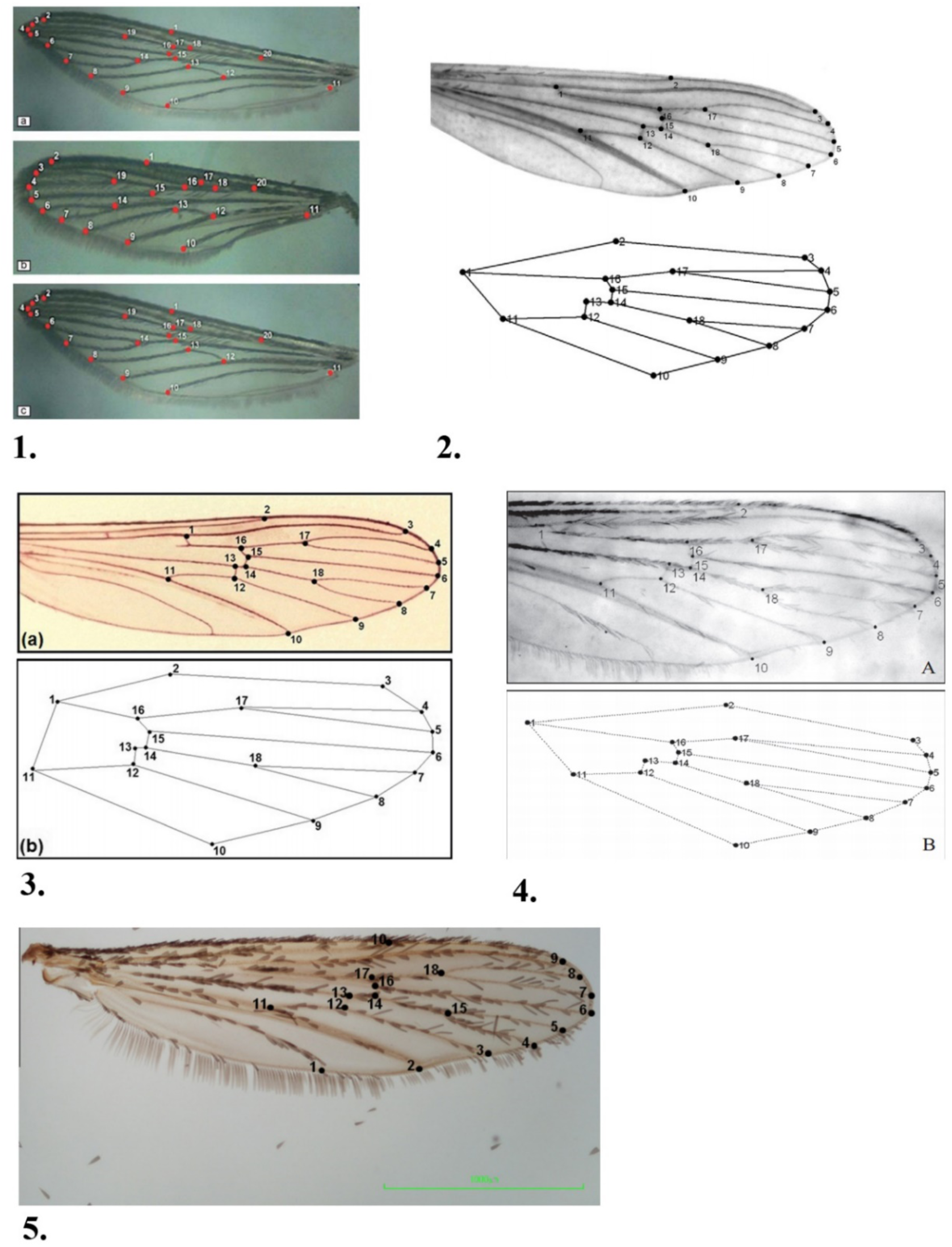
Comparison between landmark points in different mosquito wing morphometric studies. *1:* Landmarks on *Aedes aegypti* (top), *Aedes albopictus* (middle) and *Aedes pseudotaeniatus* (bottom) [50]. *2:* landmarks on *Aedes aegypti*. [52]. *3:* Landmarks on *Anopheles bellator*. [49]. *4:* Landmarks on *Culex quinquefasciatus*. [51]. *5:* Landmarks on *Aedes albopictus*. [53].

## 2. Material & Methods

### 2.1 OrchID Artificial Neural Network program

Images of mosquito wings from different genera and species were analysed by algorithms used by the OrchID [44] program. OrchID uses Artificial Neural Networks (ANN’s) to learn to recognize certain features in images. The features that can be recognized can vary by implementing different object recognition algorithms. OrchID is a Linux based program and has been used in the past to identify slipper orchids [44] and a Naturalis museum collection of Javanese butterflies (Naturalis BSc project Saskia de Vetter 2016). The orchids were classified with the help of the BGR algorithm. The BGR (Blue-Green-Red) method divided an image into horizontal and vertical bins and then calculated the mean blue-green-red values for each bin. These values were then used as features for image comparison. Using this method, slipper orchid classification accuracy was 75% for the genus level and 48% for the species level. The BGR method was also used for the Javanese butterflies, along with a second algorithm; the SURF method. The SURF (Speeded Up Robust Features) algorithm looks for changes in pixel intensity in an image to find features. The Bag-Of-Words (BOW) algorithm was then used to cluster these features for further image comparison. Using only the BGR method, butterfly genus-species accuracy was around 65%. The SURF-BOW method yielded results that were 71% accurate on genus-species level. A combination of both methods resulted in 77% accuracy. The SURF-BOW method was however not effective on the orchid collection (25% accuracy on genus-section-species level), nor was the combination of both BGR and SURF-BOW (28% accuracy). Because of these differences, the mosquito wing collection was tested with BGR, SURF-BOW and a combination of the two to find the most accurate results.

### 2.2 Image database creation

Photographs of mosquito wings were uploaded to the website “Flickr” (http://www.flickr.com) using an account created to store orchid and butterfly photographs from previous research. All mosquito wing photographs were uploaded to the folder “mosquito wings.” Flickr was used to store the photographs because the website allows users to add metadata to the pictures and because it is accessible by an application-programming interface (api), allowing for easy retrieval of the photographs along with their metadata. Metadata was included by the use of “tags.” The tags “genus:<*name*>,” “species:<*name*>,” “sex:<*name*>” and “project:mosquitoes” were given to each photograph. A small set of 37 mosquito wings was photographed with two different cameras. These photographs were given the tag “phone:S5” or “phone:G4,” depending on which camera they were photographed with. A part of the dataset was collected during fieldwork in South Africa. These photographs were given the tag “location:SA.”

#### Preparing and photographing

Because the ultimate goal of this project is to create an app that will be usable by anyone on a smartphone or tablet, all photographs of mosquito wings have been made with a smartphone. My personal smartphone, a Samsung Galaxy S5 Neo, was used as the main device for this project. A subset of photographs was taken with a Motorola Moto G4 Plus, provided by Dr. Maarten Schrama, to see how the identification program reacts to different cameras. Mosquito wings are on average only a few millimetres long and cannot be photographed in detail with the cameras used in current smartphones. A clip-on macro lens attachment was used to increase detail and reduce focusing distance. The attachment (“Clip on lens set for smartphones,” Hema) was used for all photographs. Macro lens attachments for smartphones are cheap and readily available both in physical shops and online. It is therefore presumed that the need for this tool will not diminish the functionality of a potential future app.

Mosquitoes were placed under a stereoscope and had their right wing removed using tweezers. Wings were then placed on a white sheet of paper with a colour calibration chart and a 5-mm scale bar printed on it (Figure 2). The smartphone with lens attachment was suspended 2 cm above the mosquito wing. Initially this was done by placing the phone on two Styrofoam blocks. Later two sponges were used instead of the Styrofoam. The camera function of the phone was selected and the camera was focused on the wing by tapping the phone screen in the appropriate spot. As pushing the shutter button can introduce camera shake, especially when working on macro scale, a timer was used to take the photograph. Photographs were transferred to a personal laptop (Acer Aspire ES15) using a USB cable and then uploaded to the Flickr account.

**Figure 2:**
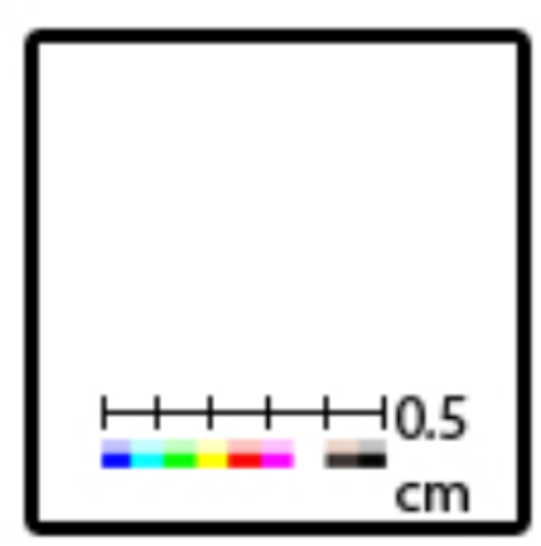
**The grid used to photograph mosquito wings on.** A small set of mosquitoes from the Naturalis collection was photographed with the wings still attached to the body. This set was used to test how accurate the image identification is if a whole (or part of a) mosquito is visible in the image. As depth of field and lighting differed greatly between individuals, depending on how the individual was positioned during preservation, the colour and scale grid was omitted from these photographs.

### 2.3 Specimen collections

Four different sets of mosquito specimen collections were used in this project. All photographed mosquitoes were either preserved in a freezer or dried. No mosquitoes preserved in liquid were used in this project. The first set was provided by Dr. Maarten Schrama and came from the Institute of Environmental Sciences. This set was mainly used as a practice and try-out set and was the only set that wasn’t photographed with the scale bar and colour calibration chart. Instead, a plain white piece of printer paper provided the background. All mosquitoes were collected and preserved the previous year. *Culex pipiens* individuals were caught at Hortus Botanicus, Leiden. All other individuals were caught in South-Africa. Identification of South-African specimens was not complete and the species of some individuals may not have been classified accurately. All genera however were accurate.

The second set of mosquitoes was provided by Wageningen University. The mosquitoes were raised in a laboratory setting. How long these mosquitoes had been preserved is unknown.

The third set came from Naturalis Biodiversity Center. This was a museum collection consisting of pinned individuals that had been preserved for many years.

The fourth set was collected during fieldwork in Kruger National Park, South-Africa, in the period March – May 2017. One part of the collection was photographed on location and one part was taken back and photographed in The Netherlands.

The total collection of photographed mosquitoes consisted of the following individuals:

#### Statistical analyses

The collections of features extracted from the images by the OrchID program algorithms were saved as text files in Tab Separated Values (tsv) format on a remote server. These files were uploaded to the online repository GitHub (www.github.com) for storage and downloaded onto a personal laptop, Acer Aspire ES15 for analysis. Analysis was done with R version 3.3.1 [56] and Rstudio version 1.0.136 [57]. The R package “ggplot2” was used for the creation of plots.

#### OrchID analyses

Initial test runs of the BGR and SURF-BOW algorithms were performed with small datasets on a personal laptop, Acer Aspire ES15. All needed programs and modules to run the tests were installed by Saskia de Vetter. Commands were run using Python 2.7. The personal laptop was not able to run the Bag Of Words algorithm on extracted features from a large dataset due to memory limitations. Final analyses with the complete dataset were therefore performed on a remote server with greater operating speed and working memory. This server was accessed using the program PuTTY Suite version 0.67. The BGR algorithm was set to divide images into 50 vertical and 50 horizontal bins for colour analysis. The Bag Of Words algorithm was set to create 100 clusters of extracted features.

## 3. Results

Results of the BGR and SURF analyses were tested with Principal Component Analyses (PCA) to visually inspect potential clustering of morphological traits. PCA’s were performed as specified by Wim van Tongeren (Naturalis MSc project 2016) and specific commands per test run can be found in Appendix III. As the number of male specimens was low, photographs of male mosquito wings were removed from analyses. Initial PCA’s showed clustering of photographs taken without a scale bar (the complete set from The Institute of Environmental Sciences, see table in appendix II), so these were also removed. A total of 475 photographs remained for final analyses.

### 3.1 BGR analysis

The 475 photographs were uploaded to the remote server and were then analysed by the BGR algorithm. The resulting datafile was downloaded onto the personal laptop and loaded into R for PCA analysis. Because one of the columns of the datafile had a constant value for all photographs, scaling was not possible. Plots were made to inspect for clustering on both the genus (figure 3) level and the species (figure 4) level. As the first two principal components explain most of the variance in the data, these parameters were used to create the x and y axes of the plots.

**Figure 3:**
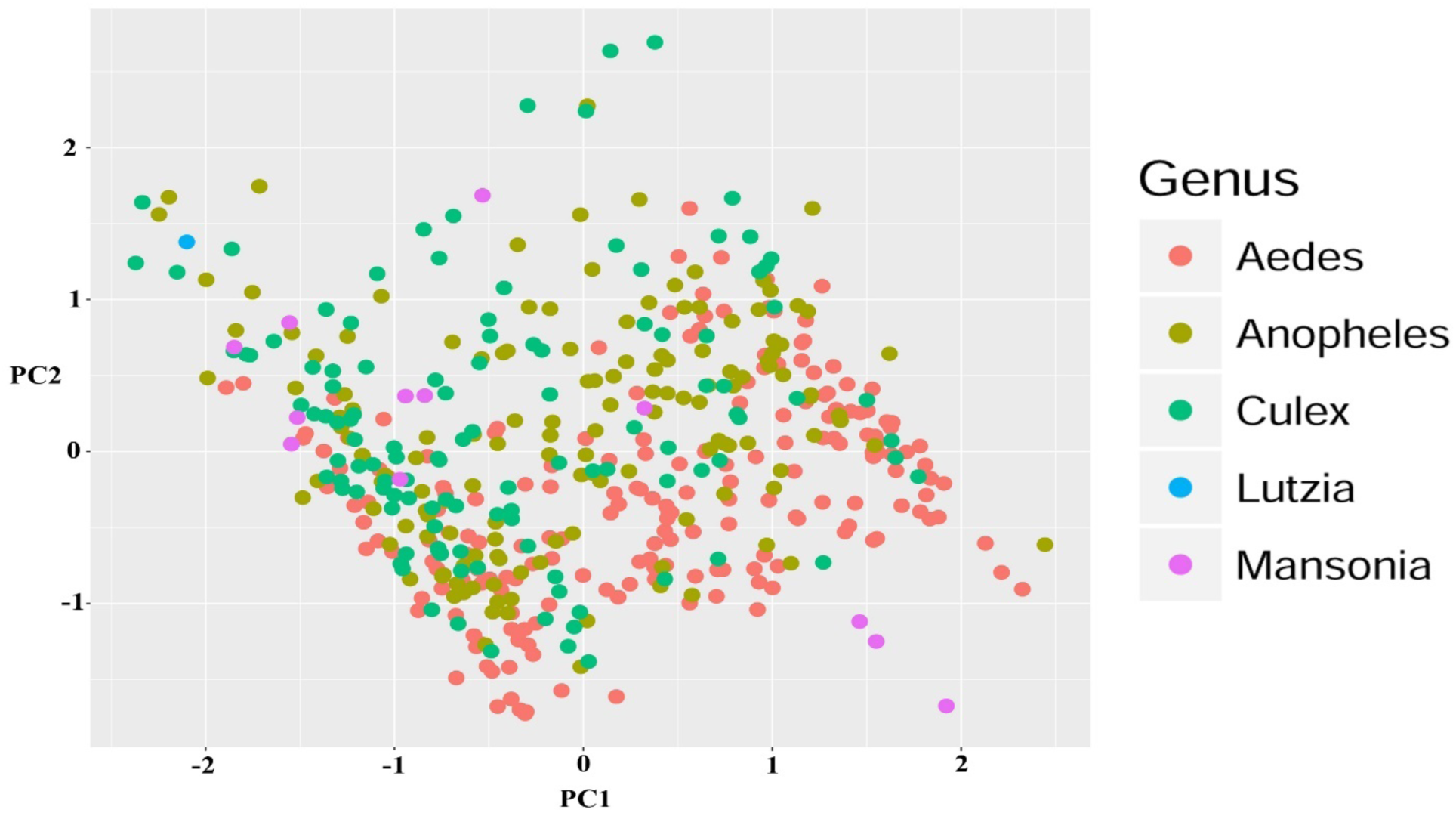
Principal component analysis plot of Blue-Green-Red algorithm analysis on mosquito wing dataset. Coloured by genus. First principal component used as x-axis, second principal component used as y-axis.

**Figure 4:**
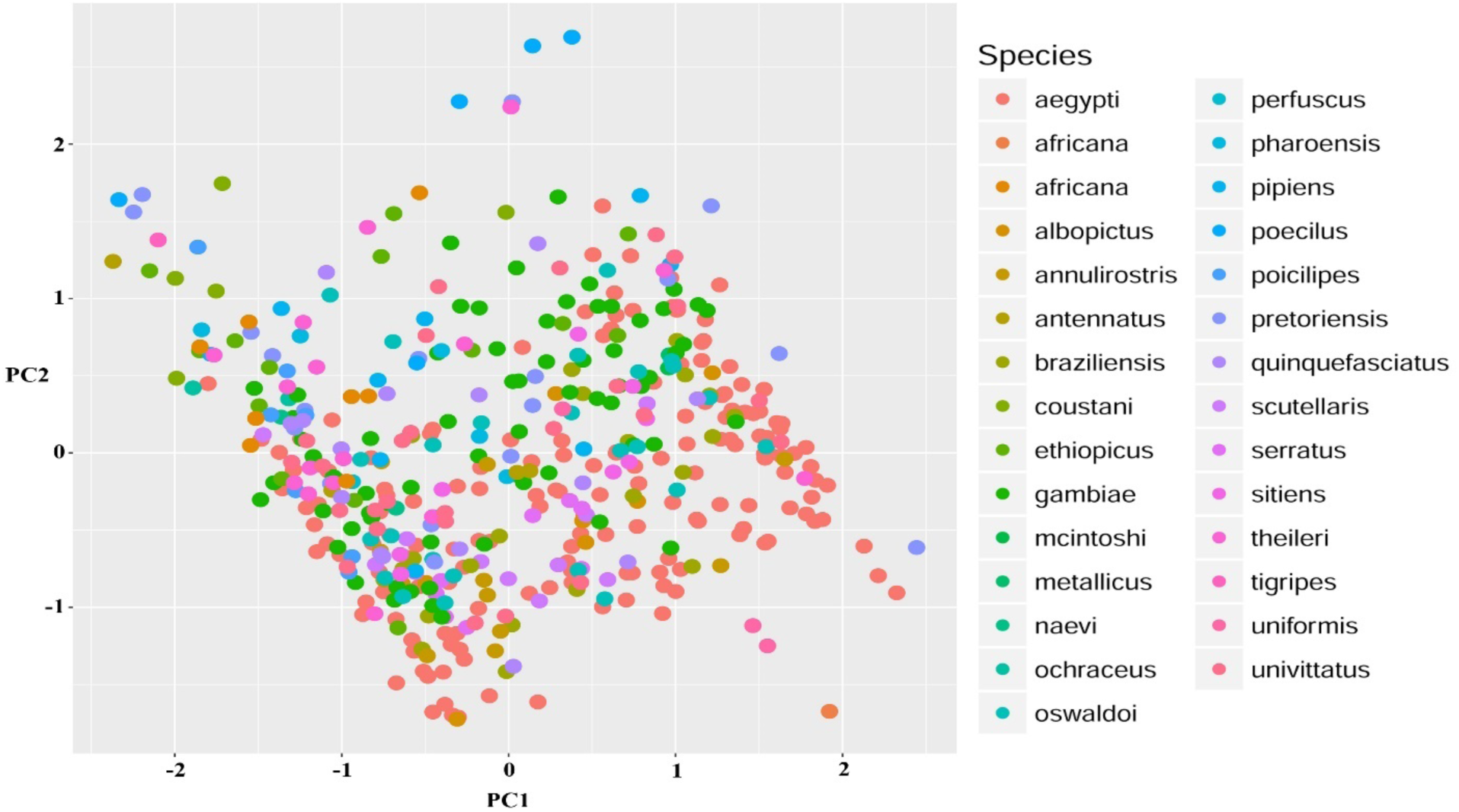
Principal component analysis plot of Blue-Green-Red algorithm analysis on mosquito wing dataset. Coloured by species. First principal component used as x-axis, second principal component used as y-axis.

### 3.2 SURF analysis

The remote server was used to analyse 475 mosquito wing photographs with the SURF algorithm. The resulting descriptions dictionary file was processed by the Bag-Of-Words algorithm which produced a dataset of clustered features for analysis in R. PCA plots were made of this dataset that visualized feature similarities on both genus (figure 5) and species (figure 6) level. The procedure was kept identical to the PCA analyses of the BGR dataset: no scaling was used and principal component 1 was used as x-axis and principal component 2 as y-axis.

**Figure 5:**
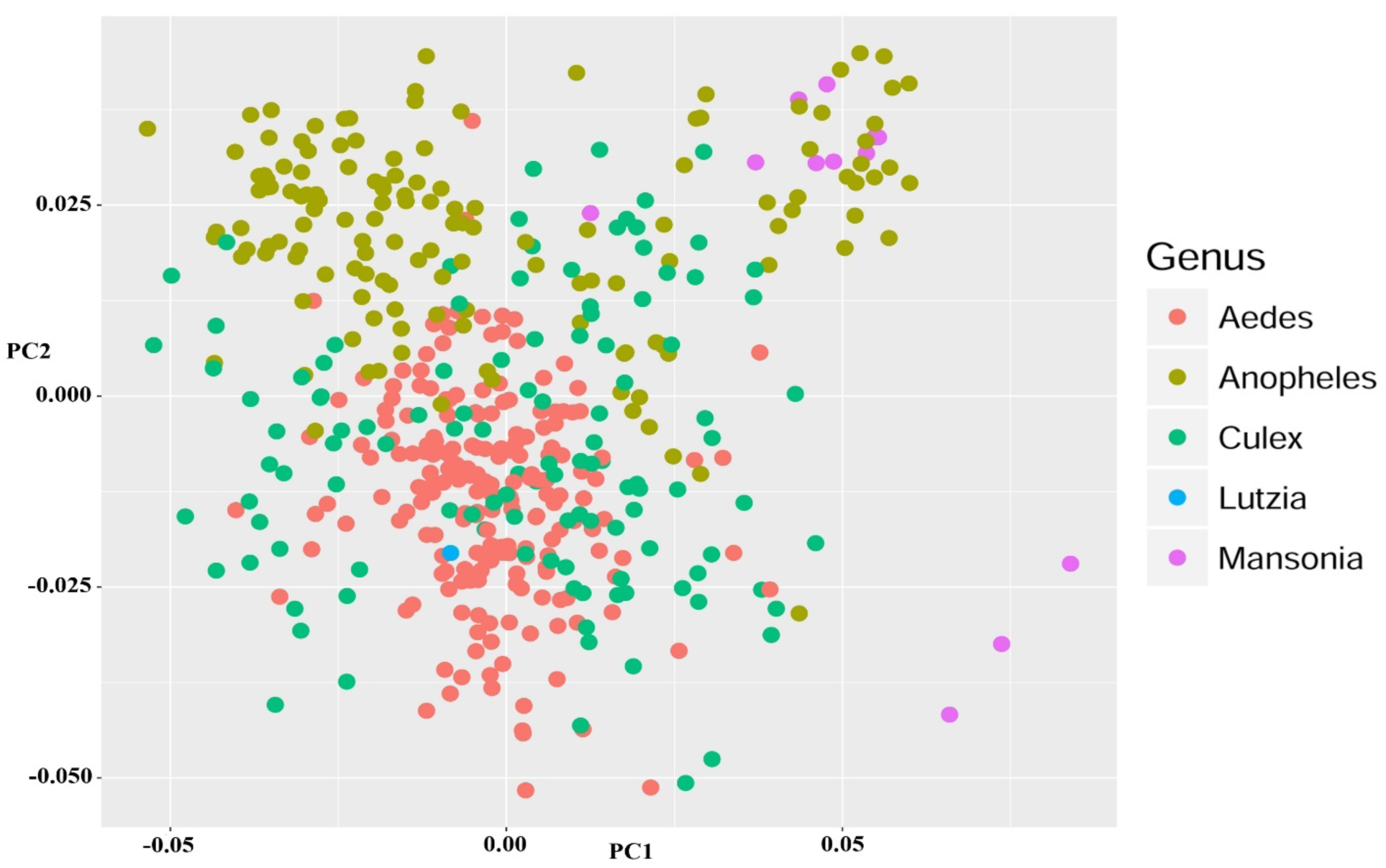
Principal component analysis plot of Speeded-Up-Robust-Features algorithm analysis on mosquito wing dataset. Coloured by genus. First principal component used as x-axis, second principal component used as y-axis.

**Figure 6:**
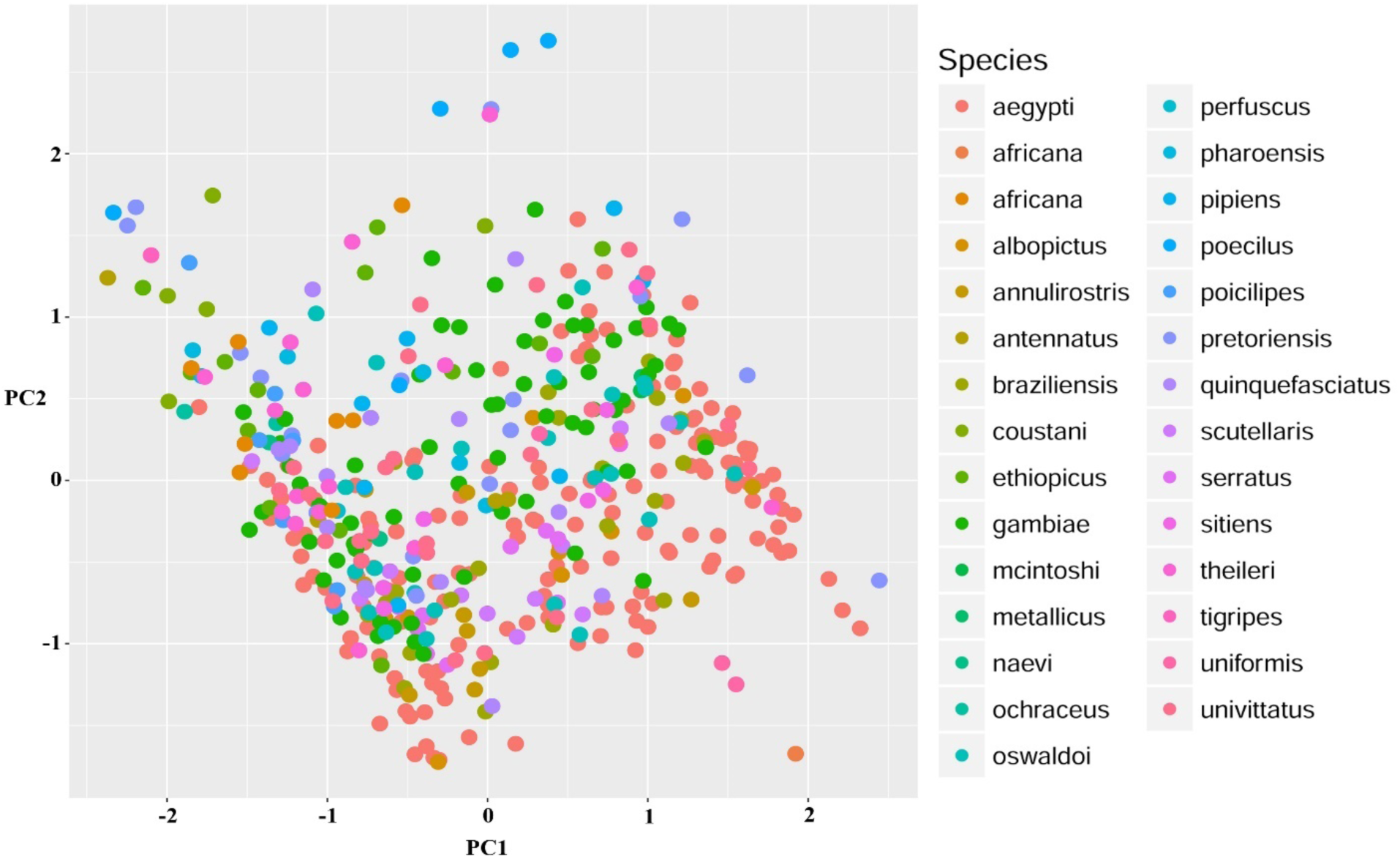
Principal component analysis plot of Speeded-Up-Robust-Features algorithm analysis on mosquito wing dataset. Coloured by species. First principal component used as x-axis, second principal component used as y-axis.

### 3.3 Phone differences

37 individuals of *Aedes aegypti* were photographed with two different smartphone cameras (Samsung S5 and Motorola G4). This subset was analysed with both the BGR (figure 7) and the SURF (figure 8) algorithms. PCA plots were created to visually inspect if the same subjects were analysed differently if photographed with a different camera.

**Figure 7:**
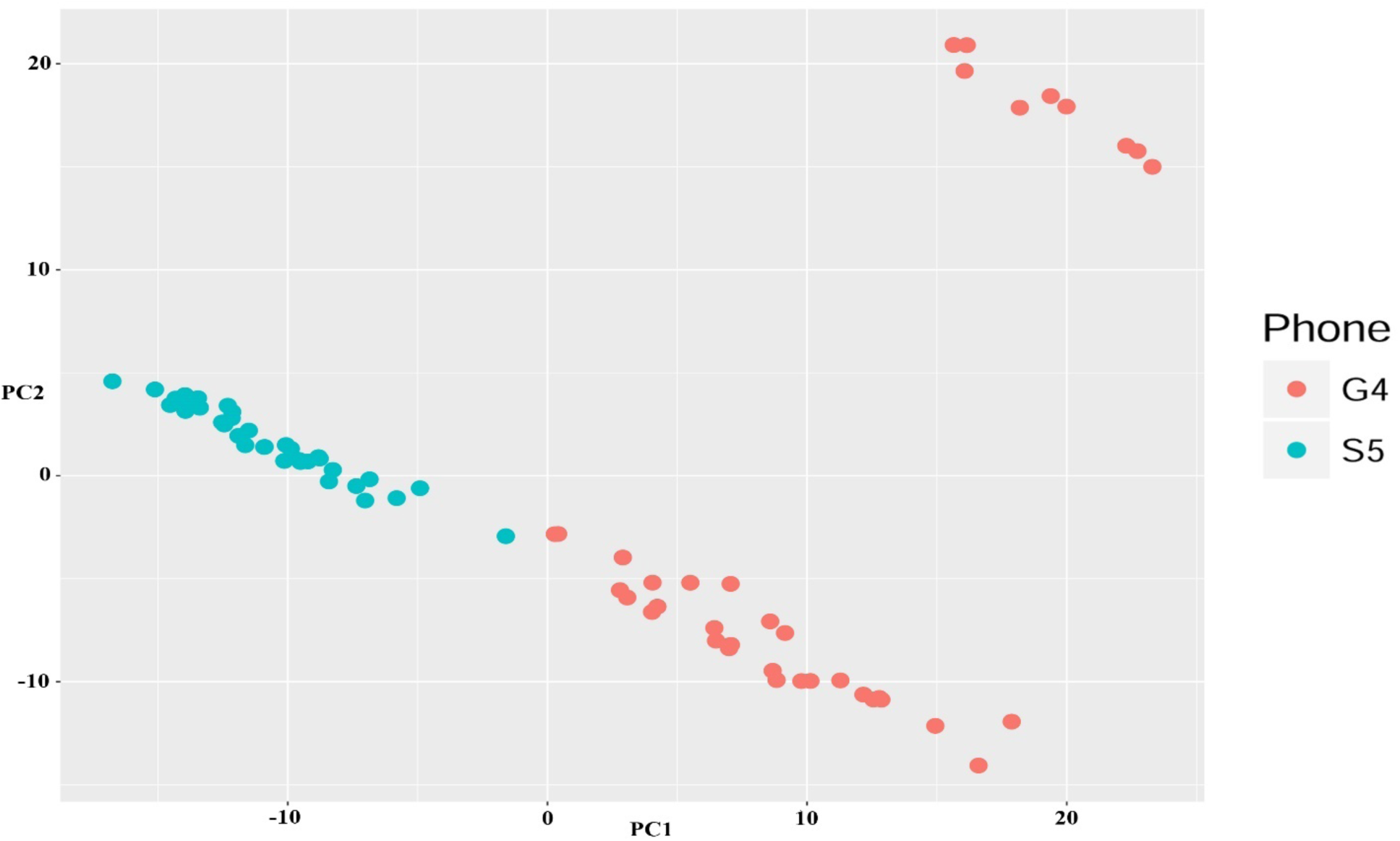
Principal component analysis plot of Blue-Green-Red algorithm analysis on 37 *Aedes aegypti* wings. Coloured by smartphone camera. First principal component used as x-axis, second principal component used as y-axis.

**Figure 8:**
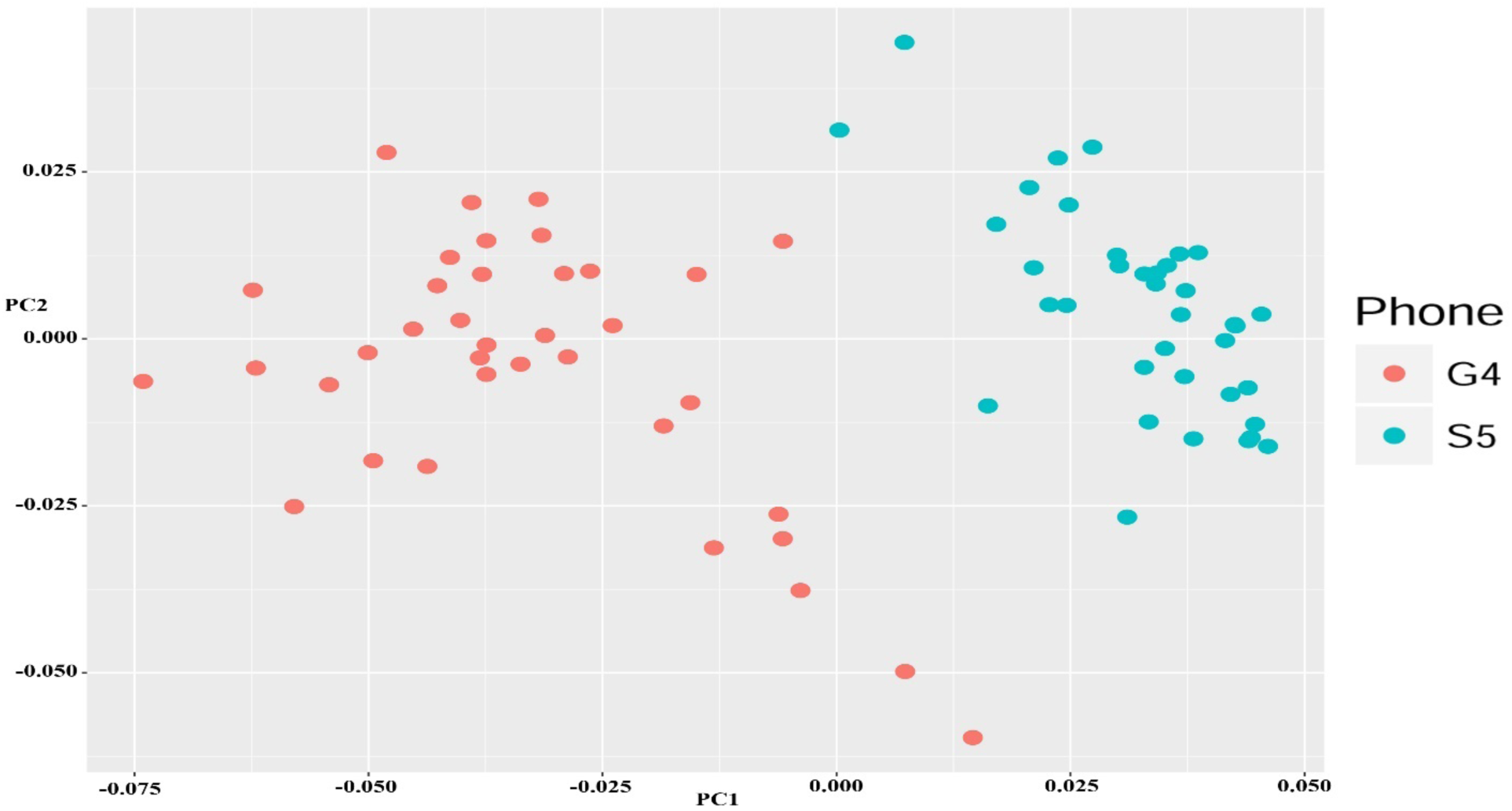
Principal component analysis plot of Speeded-Up-Robust-Features algorithm analysis on 37 *Aedes aegypti* wings. Coloured by smartphone camera. First principal component used as x-axis, second principal component used as y-axis.

## 4. Discussion

### 4.1 BGR and SURF

PCA plots of neither BGR nor SURF analyses showed clear clustering of taxonomic groups, although genera appeared somewhat more clustered than species. The two datasets were also combined so that a PCA plot could be made of both algorithms together. This however did not improve results. The BGR algorithm was programmed to divide photographs into 50 horizontal and 50 vertical bins and the Bag-Of-Words algorithm was programmed to create 100 clusters out of the SURF dataset. Tweaking these settings may provide more accurate clustering of groups. Removal of the included scale bar and colour gradient could also make for a more accurate analysis for these particular algorithms. The BGR and SURF algorithms were run with a Region of Interest (ROI) in an attempt to test this hypothesis, but the region had to be defined by width and height in pixels and the chosen region was not shown visually. This made it impossible to check if all of the scale bar and colour gradient were removed while keeping all of the wing still in the frame, as there was some variation in positioning per photograph. Because of this, ROI was left out of final analyses. In previous research, BGR proved to be more useful than SURF in orchid identification, but SURF was more effective in butterfly identification. SURF also appeared to be slightly better than BGR with the mosquito wing dataset, at least on genus level. Mosquito wings seem to differ mostly in size and shape and in areas of high contrast (black and white spots or scales), making an algorithm that analyses pixel intensity more potentially more effective than one that analyses colours.

### 4.2 Photograph examination

Though the chosen algorithms did not create clear, separate clusters of taxonomic groups, visual examination of the taken photographs show several morphological differences. The three genera comprising most of the dataset can be clearly distinguished by several features: all photographed *Anopheles* species exhibit a spotted or banded pattern created by areas of black scales (Figure 9), which are absent in *Aedes* and *Culex.* The latter two may however be dinstinguished by their shape: the wings of most photographed *Culex* species appear to be more rounded (Figure 11), whereas *Aedes* wings appear thinner and more straight (Figure 10). Differences between species are less obvious, but some species may have a great difference in wing size compared to another species in the same genus (Figures 9, 10, 11). Measuring wing size as a means for identification can however be tricky, as wing size may vary greatly even within the same species (Figure 12) and can be influenced by climate [53] and food availability [58].

**Figure 9:**
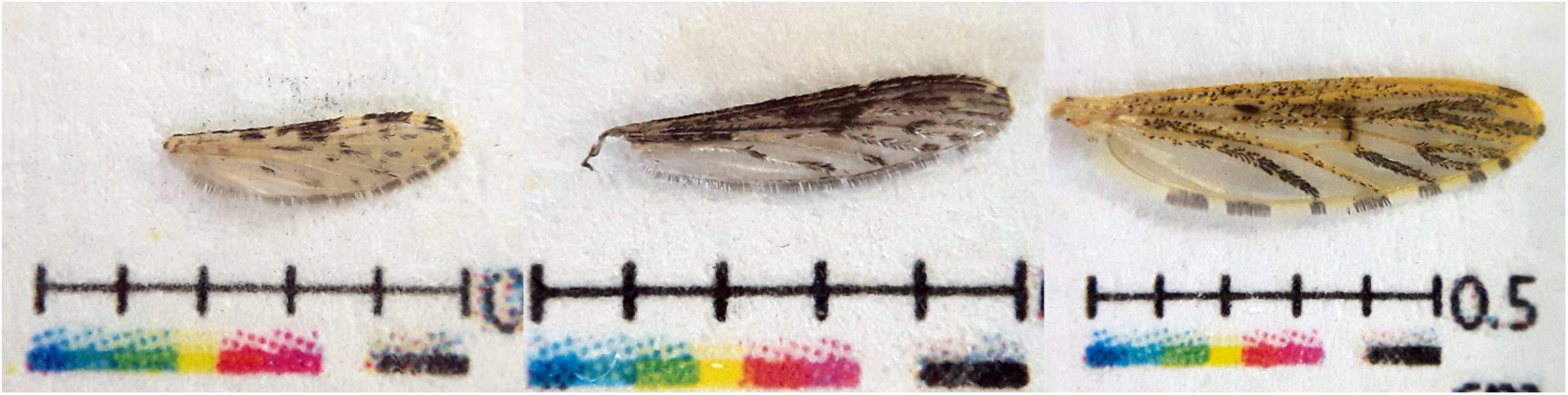
Three species of *Anopheles*. *An. gambiae* (left), *An. coustani* (middle) and *An. pharoensis* all exhibit a spotted or banded black pattern. Wing sizes differ between the photographs and may be an identifying feature between species.

**Figure 10:**
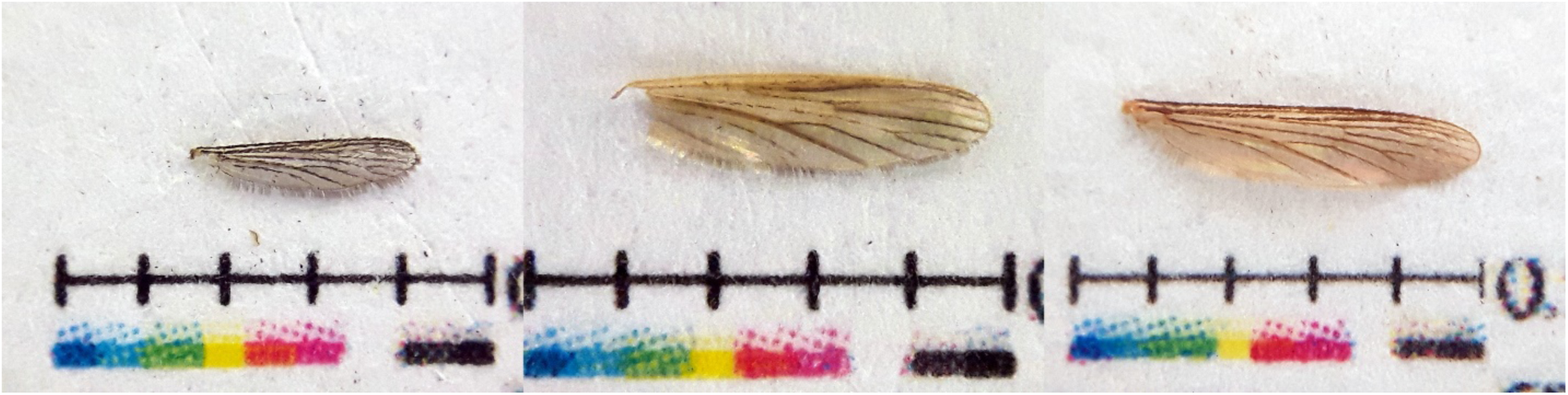
Three species of *Aedes. Ae. aegypti, Ae. ochraceus* and *Ae. scutellaris* all have relatively straight and thin wings. Wing sizes differ between the photographs and may be an identifying feature between species.

**Figure 11:**
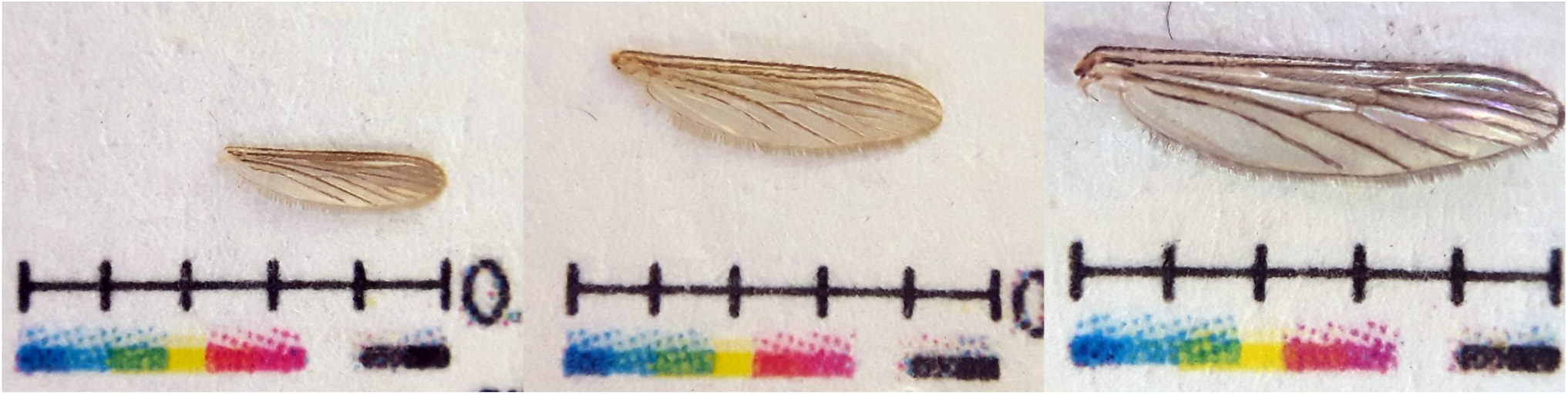
Three species of *Culex. Cu. sitiens, Cu. quinquefasciatus and Cu. theileri* wings all have relatively rounded bottom edges. Wing sizes differ between the photographs and may be an identifying feature between species.

**Figure 12:**
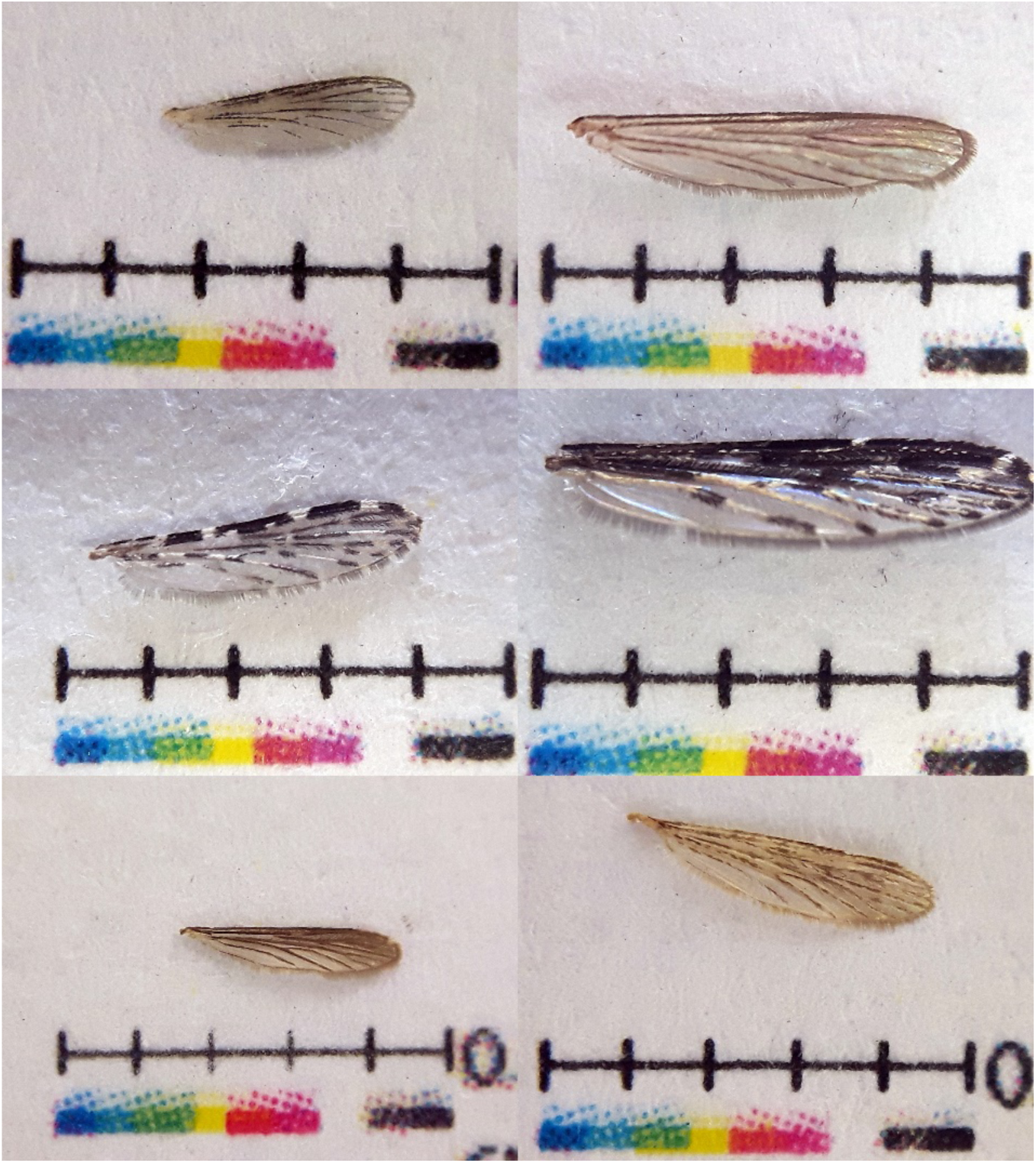
Variance in wing size within species. Top: *Culex univittatus,* Middle: *Anopheles pretoriensis,* bottom: *Aedes scutellaris.*

**Figure 13:**
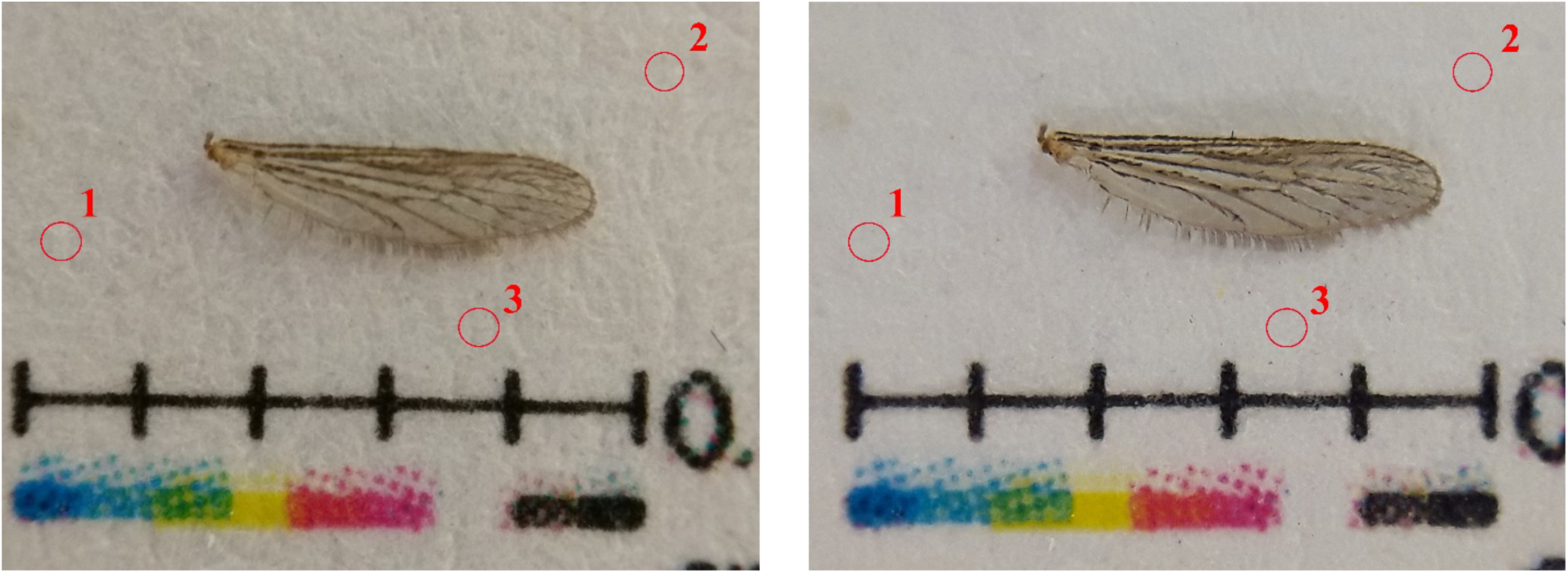
Comparison between two photographs of the same mosquito wing. Photo on the left was taken with Motorola G4, photo on the right with Samsung S5. The three points indicate where hue, saturation, brightness and colour balance were sampled.

### 4.3 Phone differences

PCA plots of the comparison between Samsung S5 and Motorola G4 smartphone cameras show that there is quite some discrepancy in found features. The two sets are clearly clustered in both the BGR and the SURF results and there is no overlap between points. The different results from the two smartphones can indicate that some hurdles may need to be overcome if this method of analysis is to be used to create a universal app. Upon visual inspection (Figure 12), differences between photographs of the same specimen can be clearly seen: the photograph taken with the Motorola G4 camera appears more blurred than the one taken with the Samsung S5 camera. This indicates that not all phones may be able to focus effectively at the minimal distances needed for macro photography. Although taken on the same day, at the same time and with the same artificial light, an inspection with the pipette tool in Adobe Photoshop CS2 of three randomly chosen points of the background of the picture also shows a difference in overall hue, saturation, brightness and colour balance (Table 3). Differing colour values can influence the BGR algorithm and differing levels of saturation and brightness can cause the SURF algorithm to find different levels of pixel intensity. Camera make and the type of processor in the device may contribute to slight visual differences that can cause difficulties for image recognition software.

**Table 3:**
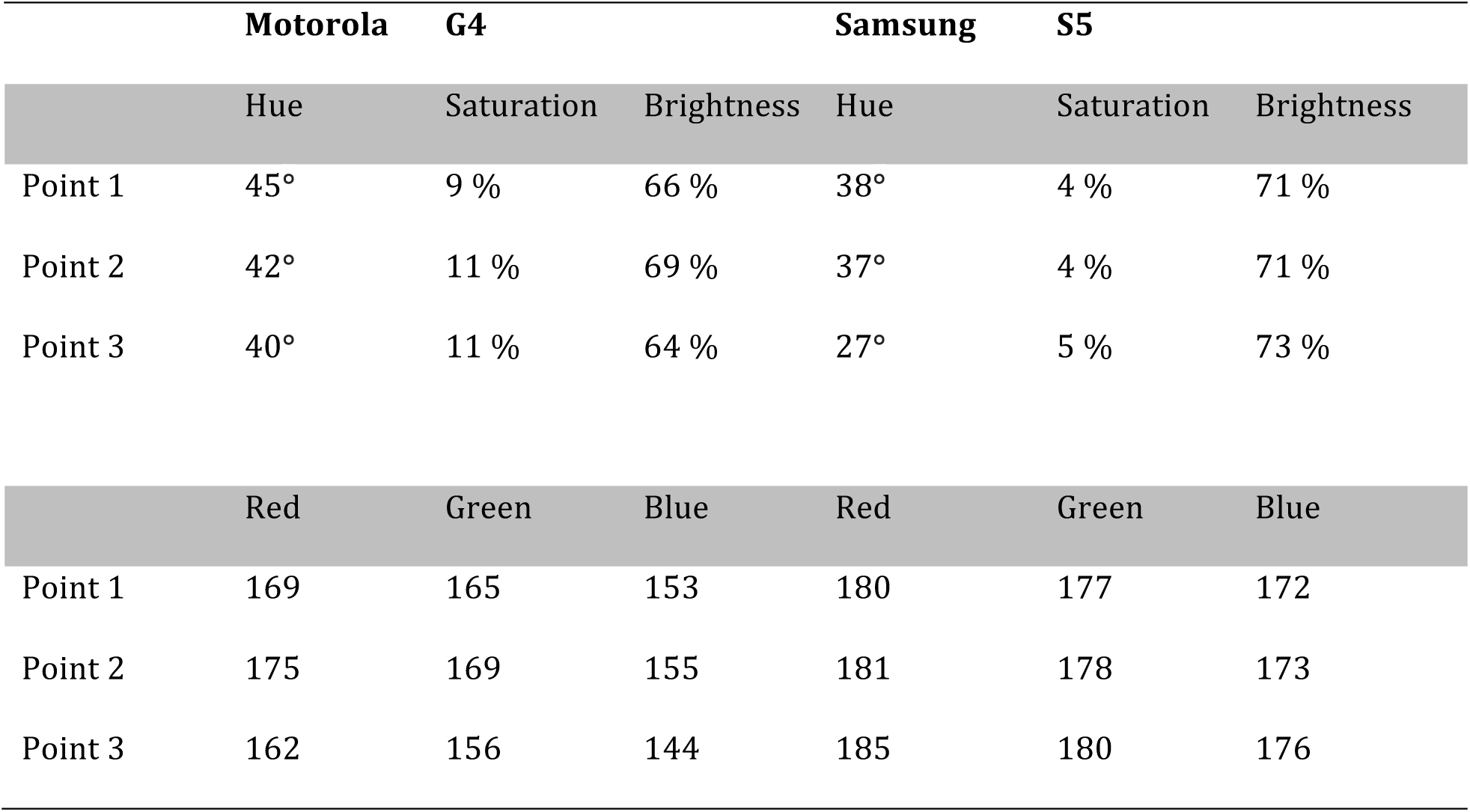
Comparison of Hue, saturation, brightness and colour balance values between Motorola G4 and Samsung S5.

|  | Motorola G4 |  |  | Samsung S5 |  |  |
| --- | --- | --- | --- | --- | --- | --- |
|  | Hue | Saturation | Brightness | Hue | Saturation | Brightness |
| Point 1 | 45° | 9 % | 66 % | 38° | 4 % | 71 % |
| Point 2 | 42° | 11 % | 69 % | 37° | 4 % | 71 % |
| Point 3 | 40° | 11 % | 64 % | 27° | 5 % | 73 % |
|  | Red | Green | Blue | Red | Green | Blue |
| Point 1 | 169 | 165 | 153 | 180 | 177 | 172 |
| Point 2 | 175 | 169 | 155 | 181 | 178 | 173 |
| Point 3 | 162 | 156 | 144 | 185 | 180 | 176 |

### 4.3 OrchID as an app

The aim of this study was to 1) test if mosquito wings are morphologically distinct enough to allow image recognition software to recognize features unique to a taxon, and 2) take the first steps in investigating if and how the OrchID program may be used as a universal identification application that would be downloadable on mobile devices. Although BGR and SURF results were far from perfect, some clustering did become apparent, especially on the genus level for the SURF algorithm. This shows that taxonomic groups of mosquitoes do indeed possess unique qualities recognizable by image recognition software. The study with landmark-based measurements by Wilke et al. (2016) also shows clear differences between genera in wing shape and size. The two algorithms used in this study did not measure these two parameters, but additional algorithms could be added to the OrchID program to make the automated identification process more effective. The implementation of the scale bar in this study could be part of a method of standardized wing photography that would allow a machine learning tool to measure different dimensions within a photograph. The added colour gradient may also be used to equalize colour levels for all photographs to make the BGR algorithm more efficient, something that has not been done in the current study. By refining and standardizing mosquito wing photography and by adding additional algorithms that would allow the program to make automated measurements of structures in a photograph, OrchID may become more precise at identifying taxonomic mosquito groups based on wing features.

## Appendix I

**Table 1:**
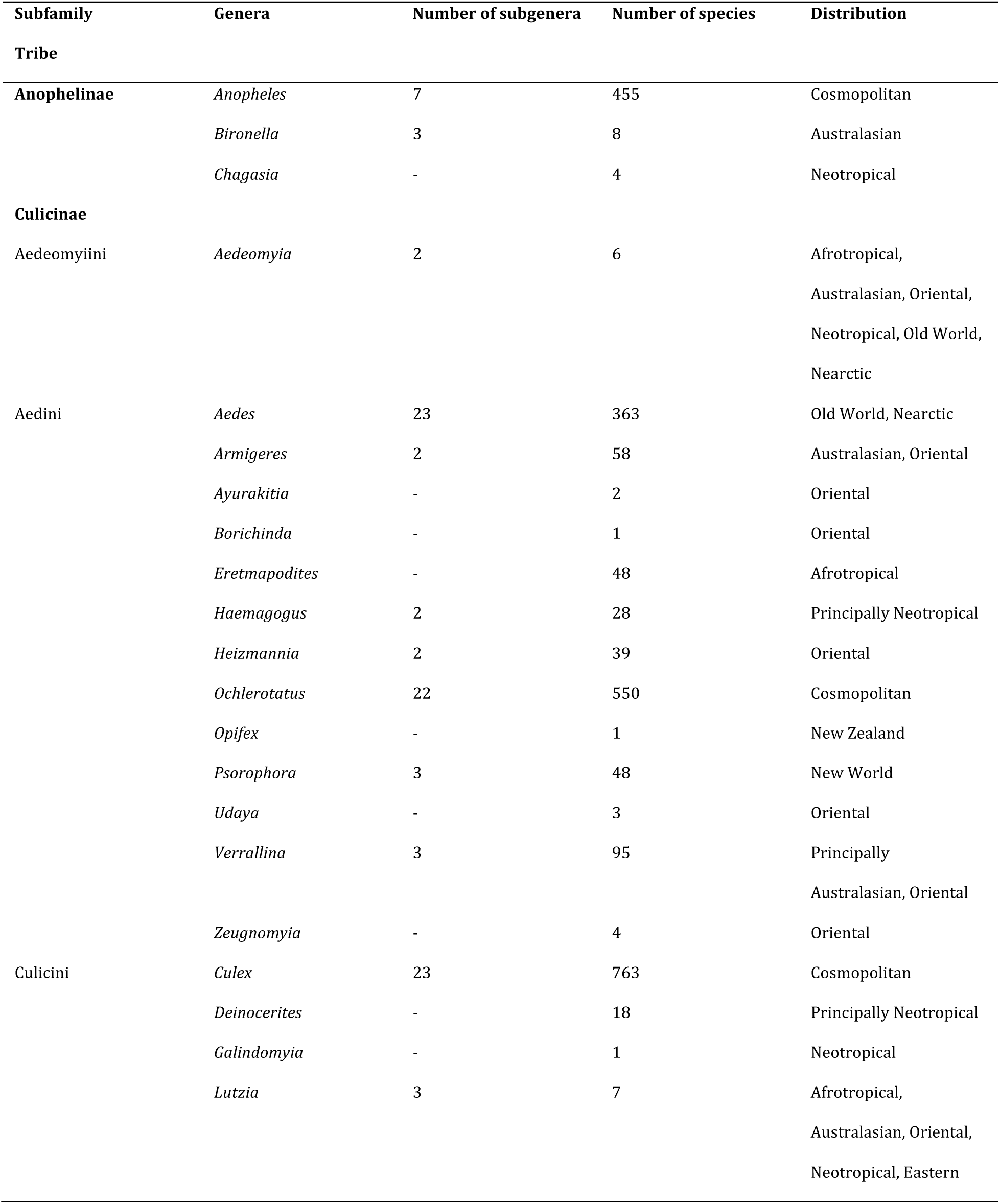
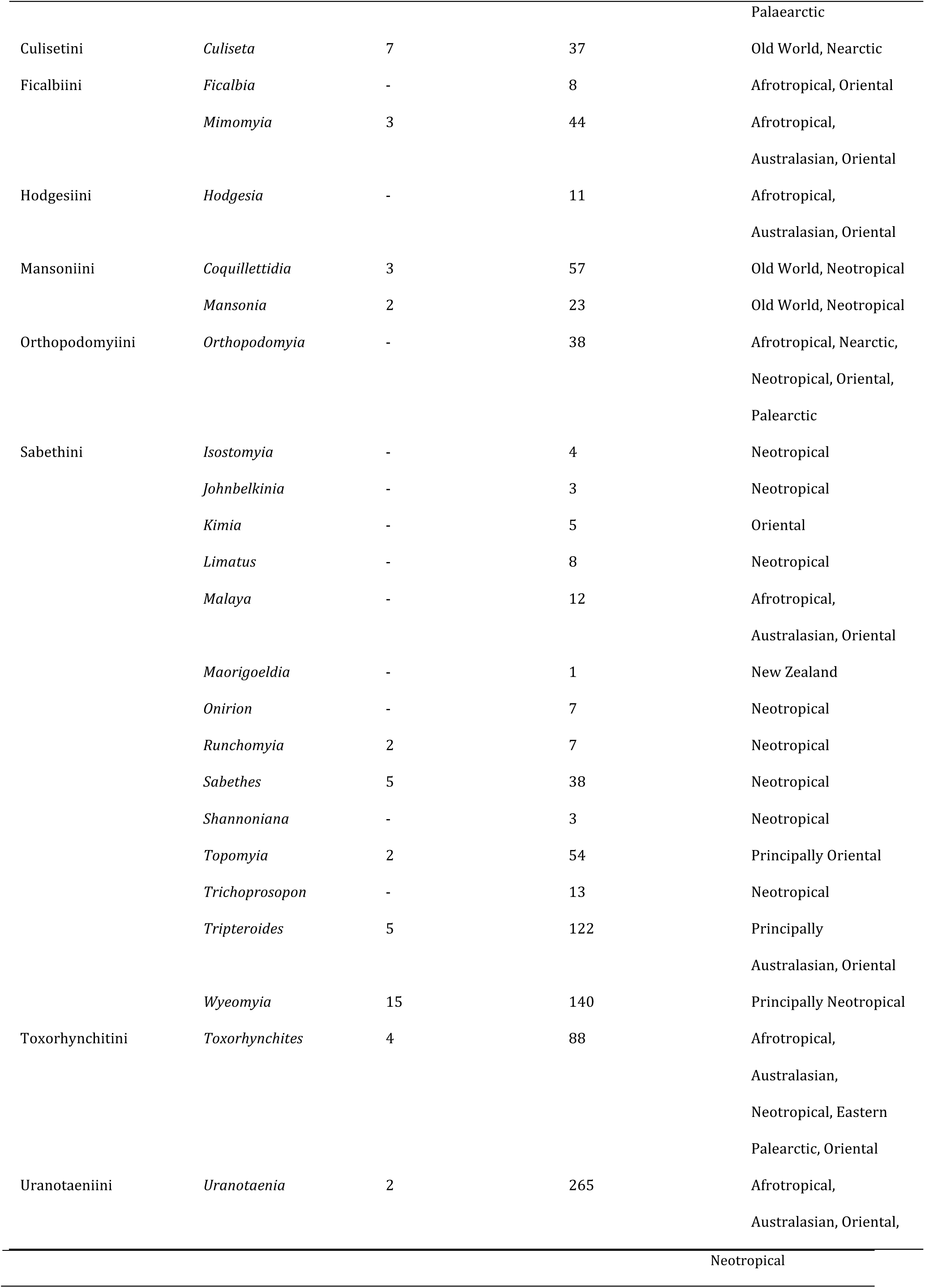
Current taxonomy and distribution of Culicidae. Taken from Harbach (2007)

## Appendix II

**Table 2:** Mosquito collection from Institute of Environmental Sciences

| Genus | Species | Number |
| --- | --- | --- |
| *Aedes* | *aegypti* | 2 females |
| *Aedes* | *albopictus* | 1 female |
| *Aedes* | Unknown | 3 females |
| *Anopheles* | Unknown | 6 females |
| *Culex* | *annuloris* | 3 females |
| *Culex* | *antennatus* | 5 females |
| *Culex* | *argentatus* | 3 females |
| *Culex* | *naevi* | 2 females |
| *Culex* | *perfuscus* | 3 females |
| *Culex* | *pipiens* | 18 males |
|  |  | 18 females |
| *Culex* | *poicilipes* | 2 females |
| *Culex* | *quinquefasciatus* | 4 females |
| *Culex* | *theileri* | 2 females |
| *Culex* | *univittatus* | 5 females |
| *Mansonia* | *africana* | 1 female |
| *Mansonia* | *uniformis* | 2 females |

**Table 3:** Mosquito collection from Wageningen Universiy

| Genus | Species | Number |
| --- | --- | --- |
| *Aedes* | *aegypti* | 167 females |
|  |  | 15 males |
| *Anopheles* | *Gambiae* | 67 females |
|  |  | 36 males |

**Table 4:** Mosquito collection from Naturalis Biodiversity Center

| Genus | Species | Number |
| --- | --- | --- |
| *Aedes* | *albopictus* | 15 males |
|  |  | 15 females |
| *Aedes* | *scutellaris* | 2 males |
|  |  | 10 females |
| *Aedes* | *serratus* | 9 males |
|  |  | 15 females |
| *Anopheles* | *braziliensis* | 22 females |
| *Anopheles* | *oswaldoi* | 25 females |
| *Culex* | *annulirostris* | 8 males |
|  |  | 17 females |
| *Culex* | *quinquefasciatus* | 9 males |
|  |  | 14 females |
| *Culex* | *sitiens* | 8 females |

**Table 5:** Mosquito collection from fieldwork in South Africa.

| Genus | Species | Number |
| --- | --- | --- |
| *Aedes* | *aegypti* | 1 female |
| *Aedes* | *mcintoshi* | 1 female |
| *Aedes* | *metallicus* | 1 female |
| *Aedes* | *ochraceus* | 2 females |
| *Anopheles* | *coustani* | 5 females |
| *Anopheles* | *pharoensis* | 5 females |
| *Anopheles* | *pretoriensis* | 17 females |
| *Anopheles* | *rufipes* | 1 female |
| *Culex* | *antennatus* | 3 females |
| *Culex* | *duttoni* | 1 female |
| *Culex* | *ethiopicus* | 14 females |
| *Culex* | *naevi* | 2 females |
| *Culex* | *perfuscus* | 1 female |
| *Culex* | *pipiens* | 10 females |
| *Culex* | *poecilus* | 5 females |
| *Culex* | *poicilipes* | 6 females |
| *Culex* | *theileri* | 16 females |
| *Culex* | *trifoliatus* | 1 female |
| *Culex* | *univittatus* | 25 females |
| *Lutzia* | *tigripes* | 1 female |
| *Mansonia* | *uniformis* | 1 female |

## Appendix III

### Genus and Species BGR PCA Plot

View(BGR_Scalebar_NoROI)

attach(BGR_Scalebar_NoROI)

PCA<-prcomp(BGR_Scalebar_NoROI[,-1:-3])

summary(PCA)

PCAPlot<-data.frame(PCA$x, Genus)

library(ggplot2)

p<-ggplot(PCAPlot, aes(PC1, PC2, label=Genus))

p+geom_point(aes(colour=Genus, size=4))+ggtitle(“First two components and Genus”)

PCAPlot2<-data.frame(PCA$x, Species)

p2<-ggplot(PCAPlot2, aes(PC1, PC2, label=Species))

p+geom_point(aes(colour=Species, size=4))+ggtitle(“First two components and Species”)

### Genus and Species SURF PCA Plot

View(BagOfWords_OnlyScalebar)

attach(BagOfWords_OnlyScalebar)

PCA<-prcomp(BagOfWords_OnlyScalebar[,-1:-3])

summary(PCA)

PCAPlot<-data.frame(PCA$x, Genus)

p<-ggplot(PCAPlot, aes(PC1, PC2, label=Genus))

p+geom_point(aes(colour=Genus, size=4))+ggtitle(“First two components and Genus”) PCAPlot2<-data.frame

(PCA$x, Species)

p2<-ggplot(PCAPlot2, aes(PC1, PC2, label=Species))

p2+geom_point(aes(colour=Species, size=4))+ggtitle(“First two components and Species”)

### Genus and Species SURF+BGR PCA Plot

View(BGR_SURF)

attach(BGR_SURF)

PCA<-prcomp(BGR_SURF[,-1:-3])

summary(PCA)

PCAPlot<-data.frame(PCA$x, Genus)

library(ggplot2)

Warning message:

package ‘ggplot2’ was built under R version 3.3.2

p<-ggplot(PCAPlot, aes(PC1, PC2, label=Genus)) p+geom_point(aes(colour=Genus, size=4))

PCAPlot2<-data.frame

PCAPlot2<-data.frame(PCA$x, Species)

p2<-ggplot(PCAPlot2, aes(PC1, PC2, label=Species)) p2+geom_point(aes(colour=Species, size=4))

### Differences between two phones BGR PCA Plot

View(BGRPhones)

attach(BGRPhones)

PCA<-prcomp(BGRPhones[,-1:-3], loadings=TRUE, scale=TRUE, scores=TRUE, cor=TRUE)

PCAPlot<-data.frame(PCA$x, Phone)

dotplot<-ggplot(PCAPlot, aes(PC1, PC2, label=Phone)) dotplot+geom_point(aes(colour=Phone, size=4))+ggtitle(“First two components and Phone”)

### Differences between two phones SURF PCA Plot

View(SURFPhones)

attach(SURFPhones)

PCA<-prcomp(SURFPhones[,-1:-3])

summary(PCA)

PCAPlot<-data.frame(PCA$x, Phone)

library(ggplot2)

Find out what’s changed in ggplot2 at

http://github.com/tidyverse/ggplot2/releases.

Warning message:

package ‘ggplot2’ was built under R version 3.3.2

p<-ggplot(PCAPlot, aes(PC1, PC2, label=Phone))

p+geom_point(aes(colour=Phone, size=4))+ggtitle(“First two components and Phone type”)

## References

1. Harbach RE. The Culicidae (Diptera): A review of taxonomy, classification and phylogeny. Zootaxa. 2007;638(1668):591–638.

2. Durden, L. A. & Mullen, G. R. Medical and Veterinary Entomology. San Diego, Calif: Academic Press; 2002.

3. Takken, W., & Verhulst, N. O. Host Preferences of Blood-Feeding Mosquitoes. Annual Review of Entomology. 2013; 433–453. http://doi.org/10.1146/annurev-ento-120811-153618

4. Okudo, H., Toma, T., Sasaki, H., Higa, Y., Fujikawa, M., Miyagi, I. et al. Crab-hole mosquito, Ochlerotatus baisasi, feeding on mudskipper (Gobiidae : Oxudercinae) in the Ryukyu Islands, Japan. Journal of the American Mosquito Control Association. 2004; 20(1), 134–137.

5. American Mosquito Control Organization. Mosquito-Borne Diseases [Internet]. 2014 [2017 May 24]. Available from: http://www.mosquito.org/mosquito-borne-diseases

6. World Health Organization. Mosquito-borne diseases [Internet]. 2016 [2017 May 24] Available from: http://www.who.int/neglected_diseases/vector_ecology/mosquito-borne-diseases/en/

7. Snell, A.E. Identification keys to larval and adult female mosquitoes (Diptera: Culicidae) of New Zealand. New Zealand Journal of Zoology, 2005;3205(March): 99–110.

8. Wang, G., Li, C., Guo, X., Xing, D., Dong, Y., Wang, Z. et al. Identifying the Main Mosquito Species in China Based on DNA Barcoding. PLoS ONE 2012;7(10), 1–11. Available from: http://doi.org/10.1371/journal.pone.0047051

9. Harbach, R. E. & Kitching, I. J. Phylogeny and classification of the Culicidae (Diptera). Systematic Entomology, 1998;23(4), 327–370. Available from: http://doi.org/10.1046/j.1365-3113.1998.00072.x

10. Jupp, P. Mosquitoes of Southern Africa. Ekogilde Publishers; 1996. 156 p.

11. Walton, C., Somboon, P., O’Loughlin, S. M., Zhang, S., Harbach, R. E., Linton, Y. M. et al. Genetic diversity and molecular identification of mosquito species in the Anopheles maculatus group using the ITS2 region of rDNA. Infection, Genetics and Evolution, 2007;7(1), 93–102. Available from: http://doi.org/10.1016/j.meegid.2006.05.001

12. Crans, W. J. A classification system for mosquito life cycles: life cycle types for mosquitoes of the northeastern United States. Journal of Vector Ecology : Journal of the Society for Vector Ecology, 2004;29(1), 1–10.

13. The Biology of Mosquitoes [Internet]. n.d. [2017 May 24]. Available from: http://www.smsl.co.nz/site/southernmonitoring/images/NZB/MossieAwareness/The%20Biology%20of%20Mosquitoes.pdf

14. Mukhtar M., Herrel N., Amerasinghe F.P., Ensink J., van der Hoek W., Konradsen F. Role of wastewater irrigation in mosquito breeding in south Punjab, Pakistan. Southeast Asian Journal of Tropical Medicine and Public Health. 2003;34: 72–80.

15. Dar es Salaam Urban Malaria Control Programme. Guidelines to searching for mosquito breeding habitats (stagnant water) and conducting larval survey. 2005. Available from: http://www.biomedcentral.com/content/supplementary/1475-2875-7-20-S2.pdf

16. Mosquito World. Mosquito habitats. N.d. [2017 May 24]. Available from: http://www.mosquitoworld.net/about-mosquitoes/habitats/

17. Okogun, G. R., Nwoke, B. E., Okere, A. N., Anosike, J. C., & Esekhegbe, A. C. Epidemiological implications of preferences of breeding sites of mosquito species in Midwestern Nigeria. Annals of Agriculture and Environmental Medicine. 2003;10(2), 217–222. Available from: http://doi.org/10.1111/j.1440-1843.2003.supp_1.x

18. Gunathilaka, N., Fernando, T., Hapugoda, M., Wickremasinghe, R., & Wijeyerathne, P. Anopheles culicifacies breeding in polluted water bodies in Trincomalee District of Sri Lanka. Malaria Journal. 2013; 12(285). Available from: http://doi.org/10.1186/1475-2875-12-285

19. Kudom, A. A. Larval ecology of Anopheles coluzzii in Cape Coast, Ghana: water quality, nature of habitat and implication for larval control. Malaria Journal. 2015;14(1), 447. Available from: http://doi.org/10.1186/s12936-015-0989-4

20. De Lamballerie, X., Leroy, E., Charrel, R. N., Ttsetsarkin, K., Higgs, S., & Gould, E. A. Chikungunya virus adapts to tiger mosquito via evolutionary convergence: a sign of things to come? Virology Journal. 2008;5, 33. Available from: http://doi.org/10.1186/1743-422X-5-33

21. Lambrechts, L., Scott, T. W., & Gubler, D. J. Consequences of the expanding global distribution of Aedes albopictus for dengue virus transmission. PLoS Neglected Tropical Diseases. 2010;4(5). Available from: http://doi.org/10.1371/journal.pntd.0000646

22. Wada, Y. Japanese Encephalitis Vectors. Tropical Medicine. 1995;36(4), 235–242. Available from: http://naosite.lb.nagasaki-u.ac.jp/dspace/bitstream/10069/4695/1/tm36_04_16_t.pdf

23. Rogers, D. J., Randolph, S. E., Snow, R. W., & Hay, S. I. Satellite imagery in the study and forecast of malaria. Nature. 2002;415(6872), 710–715. http://doi.org/10.1038/415710a

24. World Health Organization. Do all mosquitoes transmit malaria [Internet]? 2016 [2017 May 24]. Available from: http://www.who.int/features/qa/10/en/

25. Hubálek, Z., & Halouzka, J. West Nile fever--a reemerging mosquito-borne viral disease in Europe. Emerging Infectious Diseases. 1999;5(5), 643–650. Available from: http://doi.org/10.3201/eid0505.990506

26. Gubler, D. J. The changing epidemiology of yellow fever and dengue, 1900 to 2003: Full circle? Comparative Immunology, Microbiology and Infectious Diseases. 2004;27(5), 319–330. Available from: http://doi.org/10.1016/j.cimid.2004.03.013

27. Diallo, D., Sall, A. A., Diagne, C. T., Faye, O., Faye, O., Ba, Y. et al. Zika virus emergence in mosquitoes in Southeastern Senegal, 2011. PLoS ONE. 2014; 9(10), 4–11. Available from: http://doi.org/10.1371/journal.pone.0109442

28. Cano, J., Rebollo, M. P., Golding, N., Pullan, R. L., Crellen, T., Soler, A. et al. The global distribution and transmission limits of lymphatic filariasis: past and present. Parasites & Vectors. 2014;7(1), 466. Available from: http://doi.org/10.1186/s13071-014-0466-x

29. World Health Organization. Lymphatic filariasis [Internet]. 2016 [2017 May 24]. Available from: http://www.who.int/lymphatic_filariasis/epidemiology/en/

30. CMETE. Diseases transmitted by insects and ticks [Internet]. N.d. [2017 May 24]. Available from: http://www.cmete.com/index.php?tg=articles&idx=Print&topics=101&article=68

31. Agyepong, I. A. Malaria: Ethnomedical perceptions and practice in an Adangbe farming community and implications for control. Social Science and Medicine. 1992;35(2), 131–137. Available from: http://doi.org/10.1016/0277-9536(92)90160-R

32. Ahorlu, C. K., Dunyo, S. K., Afari, E. A., Koram, K. A., & Nkrumah, F. K. Malaria-related beliefs and behaviour in southern Ghana: implications for treatment, prevention and control. Tropical Medicine & International Health : TM & IH. 1997;2(5), 488–499. Available form: http://doi.org/10.1111/j.1365-3156.1997.tb00172.x

33. Deressa, W., Ali, A., & Enquoselassie, F. Knowledge, Attitude and Practice About Malaria, the Mosquito and Antimalarial Drugs in a Rural Community. Ethiop. J. Health Dev. 2004;100–105.

34. Epstein, P. R., Diaz, S., Elias, T. S., Grabherr, G., Graham, N., Martens, W. et al. Biological and physical signs of climate change: focus on mosquito borne diseases. Bulletin of the American Meteorological Society. 1998;79, 405–417.

35. Bedford, G. A. South African Mosquitoes. Reports of the Director of Veterinary Education and Research. 1928;13 & 14, 883–900. Available from: http://repository.up.ac.za/bitstream/handle/2263/13672/883-900_bdf-1.pdf

36. Kumar, N. P., Rajavel, A. R., Natarajan, R., & Jambulingam, P. DNA barcodes can distinguish species of Indian mosquitoes (Diptera: Culicidae). Journal of Medical Entomology. 2007;44(1), 1–7. Available from: http://doi.org/10.1093/jmedent/41.5.01

37. Wilke, A. B., De Oliveira Christe, R., Multini, L. C., Vidal, P. O., Wilk-Da-silva, R., De Carvalho, G. C. et al. Morphometric wing characters as a tool for mosquito identification. PLoS ONE. 2016;11(8), 1–12. Available from: http://doi.org/10.1371/journal.pone.0161643

38. Bay, H., Ess, A., Tuytelaars, T., & Van Gool, L. Speeded-Up Robust Features (SURF). Computer Vision and Image Understanding. 2008;110(3), 346–359. Available from: http://doi.org/10.1016/j.cviu.2007.09.014

39. Gevers, T., & Smeulders, A. W. Color-based object recognition. Pattern Recognition. 1999;32(3), 453–464. Available from: http://doi.org/10.1016/S0031-3203(98)00036-3

40. Schiele, B., & Crowley, J. L. Recognition without correspondance using multidimensional receptive field histograms. International Journal of Computer Vision. 2000;36(1), 31–50.

41. Tumbling Dice. Daisy [Internet]. 2012 [2017 May 24]. Available from: http://www.tumblingdice.co.uk/daisy

42. Faria, F. A., Perre, P., Zucchi, R. A., Jorge, L. R., Lewinsohn, T. M., Rocha, A. et al. Automatic identification of fruit flies (Diptera: Tephritidae). Journal of Visual Communication and Image Representation. 2014;25(7), 1516–1527. Available from: http://doi.org/10.1016/j.jvcir.2014.06.014

43. De Meulemeester T. & Pereira S. WingID: Morphometrics-based identification app [Internet]. 2015 [2017 May 24]. Naturalis Biodiversity Center, Leiden, The Netherlands. Available from: http://wing-id.naturalis.nl/.

44. Pereira, S., Gravendeel, B., Wijntjes, P., & Vos, R. A. OrchID : a Generalized Framework for Taxonomic Classification of Images Using Evolved Artificial Neural Networks. 2016. Manuscript Submitted For Publication.

45. Russell, K. N., Do, M. T., Huff, J. C., & Platnick, N. I. Introducing SPIDA-Web: Wavelets, neural networks and internet accessibility in an image-based automated identification system. Automated Taxon Identification in Systematics: Theory, Approaches and Applications. 2007;74, 131–152.

46. Tofilski, A. DrawWing, a program for numerical description of insect wings. Journal of Insect Science. 2004;4(17), 1–5. Available from: http://doi.org/10.1673/031.004.1701

47. Francoy, T. M., Wittmann, D., Drauschke, M., Muller, S., Steinhage, V., Bezerra- Laure et al. Identification of Africanized honey bees through wing morphometrics: two fast and efficient procedures. Apidologie, Springer Verlag. 2008;39 (5), pp.488–494.

48. Yang, H.-P., Ma, C.-S., Wen, H., Zhan, Q.-B., & Wang, X.-L. A tool for developing an automatic insect identification system based on wing outlines. Scientific Reports 2015;5, 12786. Availablre from: http://doi.org/10.1038/srep12786

49. Lorenz, C., Marques, T., Sallum, M. A., & Suesdek, L. Morphometrical diagnosis of the malaria vectors Anopheles cruzii, An. homunculus and An. bellator. Parasites & Vectors, 2012;5(1), 257. Available from: http://doi.org/10.1186/1756-3305-5-257

50. Mondal, R., Pemola Devi, N., & Jauhari, R. K. Landmark-based geometric morphometric analysis of wing shape among certain species of Aedes mosquitoes in District Dehradun (Uttarakhand), India. Journal of Vector Borne Diseases. 2015;52(2), 122–128.

51. Vidal, P. O., Peruzin, M. C., & Suesdek, L. Wing diagnostic characters for Culex quinquefasciatus and Culex nigripalpus (Diptera, Culicidae). Revista Brasileira de Entomologia. 2011;55(1), 134–137. Available from: http://doi.org/10.1590/S0085-56262011000100022

52. Vidal, P. O., & Suesdek, L. Comparison of wing geometry data and genetic data for assessing the population structure of Aedes aegypti. Infection, Genetics and Evolution. 2011;12(3), 591–596. Available from: http://doi.org/10.1016/j.meegid.2011.11.013

53. Morales Vargas, R. E., Phumala-Morales, N., Tsunoda, T., Apiwathnasorn, C., & Dujardin, J. P. The phenetic structure of Aedes albopictus. Infection, Genetics and Evolution. 2013;13(1), 242–251. Available from: http://doi.org/10.1016/j.meegid.2012.08.008

54. Virginio, F., Oliveira Vidal, P., & Suesdek, L. Wing sexual dimorphism of pathogen-vector culicids. Parasites & Vectors. 2015;8(4), 159. Available from: http://doi.org/10.1186/s13071-015-0769-6

55. Jupp, P.G. Mosquitoes as vectors of human disease in South Africa, South African Family Practice. 2005;47(9), 68–72. Available from: doi:10.1080/20786204.2005.1087329.

56. R Core Team. R: A language and environment for statistical computing. R Foundation for 603 Statistical Computing, Vienna, Austria. 2016. Available from: https://www.R-project.org/.

57. RStudio Team. RStudio: Integrated Development for R. RStudio, Inc. Boston, MA. 2016. Available from: http://www.rstudio.com/.

58. Jirakanjanakit N, Leemingsawat S, Thongrungkiat S, Apiwathnasorn C, Singhaniyom S, Bellec C, et al. Influence of larval density or food variation on the geometry of the wing of Aedes (Stegomyia) aegypti. Trop Med Int Heal. 2007;12(11):1354–60.

